# QUAL-IF-AI: Quality Control of Immunofluorescence Images using Artificial Intelligence

**DOI:** 10.1101/2024.01.26.577391

**Authors:** Madhavi Dipak Andhari, Giulia Rinaldi, Pouya Nazari, Gautam Shankar, Nikolina Dubroja, Johanna Vets, Tessa Ostyn, Maxime Vanmechelen, Brecht Decraene, Alexandre Arnould, Willem Mestdagh, Bart De Moor, Frederik De Smet, Francesca Bosisio, Asier Antoranz

## Abstract

Fluorescent imaging has revolutionized biomedical research, enabling the study of intricate cellular processes. Multiplex immunofluorescent imaging has extended this capability, permitting the simultaneous detection of multiple markers within a single tissue section. However, these images are susceptible to a myriad of undesired artifacts, which compromise the accuracy of downstream analyses. Manual artifact removal is impractical given the large number of images generated in these experiments, necessitating automated solutions. Here, we present QUAL-IF-AI, a multi-step deep learning-based tool for automated artifact identification and management. We demonstrate the utility of QUAL-IF-AI in detecting four of the most common types of artifacts in fluorescent imaging: air bubbles, tissue folds, external artifacts, and out-of-focus areas. We show how QUAL-IF-AI outperforms state-of-the-art methodologies in a variety of multiplexing platforms achieving over 85% of classification accuracy and more than 0.6 Intersection over Union (IoU) across all artifact types. In summary, this work presents an automated, accessible, and reliable tool for artifact detection and management in fluorescent microscopy, facilitating precise analysis of multiplexed immunofluorescence images.

## Introduction

Fluorescent imaging has become an indispensable tool in various biological disciplines, enabling the study of protein dynamics, cellular functions, and neuronal activity, among others [1]. With the recent development of multiplex immunofluorescent imaging, the simultaneous detection and quantification of multiple markers in a single tissue section has become feasible [2]. The analysis of the images generated with multiplex immunofluorescent imaging requires an extensive pipeline of preprocessing steps, including illumination correction, image registration (in the case of cyclic multiplexing methods), cell segmentation and autofluorescence subtraction before extracting quantitative single-cell measurements from the images [3]–[5]. The quality of the input images is crucial to obtain reliable single-cell measurements and meaningful biological insights. Undesired artifacts, such as external artifacts (dust particles, hair shafts, fibers), air bubbles, antibody aggregates, out-of-focus (OOF) areas, and tissue folds are common not only in hematoxylin and eosin (H&E) images [6] but also in fluorescent microscopy [7]. Dust particles, hair shafts, fibers, and air bubbles can get trapped during sample preparation or coverslipping. Denatured antibodies can lead to antibody aggregation artifacts in the images. Misalignment of coverslip or inaccurate calculation of focus depth or tissue detachment during scanning can give rise to OOF areas. Suboptimal tissue embedding or tissue sectioning can lead to tissue folding or even complete tissue detachment from the slide [6]. These artifacts can significantly impact quantitative image analysis by including false positives and false negatives [8], [9]. Traditionally, artifacts are manually annotated and removed from the images. With the rise of multiplexing, the number of images generated per experiment is drastically increasing, making manual annotation extremely laborious and time-consuming. Moreover, manual annotation is subjective and contingent on high intra- and inter-user variability [10], [11]. To overcome these challenges, automated tools for detecting and managing artifacts in an unbiased and reproducible way are a necessity. Although some automated tools for detecting artifacts in H&E images exist [12]–[15], limited tools are available for fluorescent microscopy. CellProfiler [16] can identify artifacts in fluorescent microscopy, but its applicability is limited to identifying only OOF areas and saturated debris. However, air bubbles, tissue folds, and external artifacts are equally relevant and do not have suitable tools for their identification. Additionally, the sensitivity of CellProfiler to parameter adjustments introduces some degree of complexity in its usage. Therefore, the set of tools currently available in the literature falls short in terms of diversity of identifiable artifacts, ease of use, and overall performance in the context of fluorescent microscopy images.

The increase in the popularity of deep learning (DL), and particularly convolutional neural networks (CNN) for image analysis, is starting to show their applicability for artifact detection in medical Images [17], [18]. One of the main bottlenecks of these approaches is that training models with high accuracy requires pixel level annotations for thousands of images [18]. Transfer learning can partially address this issue by leveraging information learned on other bigger datasets such as ImageNet, a popular visual database containing over 14 million manually annotated images which has been used to train a variety of DL architectures. Several models have been shown to perform better with transfer learning in classifying medical images for different tasks [19]–[21]. For example, a study conducted by Guan et al. demonstrated the effectiveness of transfer learning in differentiating papillary thyroid carcinoma from benign thyroid nodules using cytological images. By fine-tuning pre trained VGG16 and Inception v3 models on relatively small datasets they achieved remarkable accuracy in identifying papillary thyroid carcinoma [21].

Building upon these ideas, we have developed QUAL-IF-AI (reads qualify), a DL-based two-tier quality control tool for the detection and management of four different types of artifacts: OOF areas, external artifacts, tissue folds, and air bubbles. QUAL-IF-AI first classifies tissue regions into having the presence or absence of given artifacts and then delineates (segments) artifacts if the initial prediction is positive. We benchmarked a series of classification (VGG16, VGG19, Inception v3 and Resnet50) [22] and segmentation (U-Net) [23] architectures and selected the best-performing model in each case (Fig 1). We also demonstrate that QUAL-IF-AI outperforms state-of-the-art methodologies in a series of datasets acquired with three different multiplexing technologies. Overall, in this work, we present a comprehensive quality control algorithm for artifact detection and management in fluorescent images.

**Fig 1.**
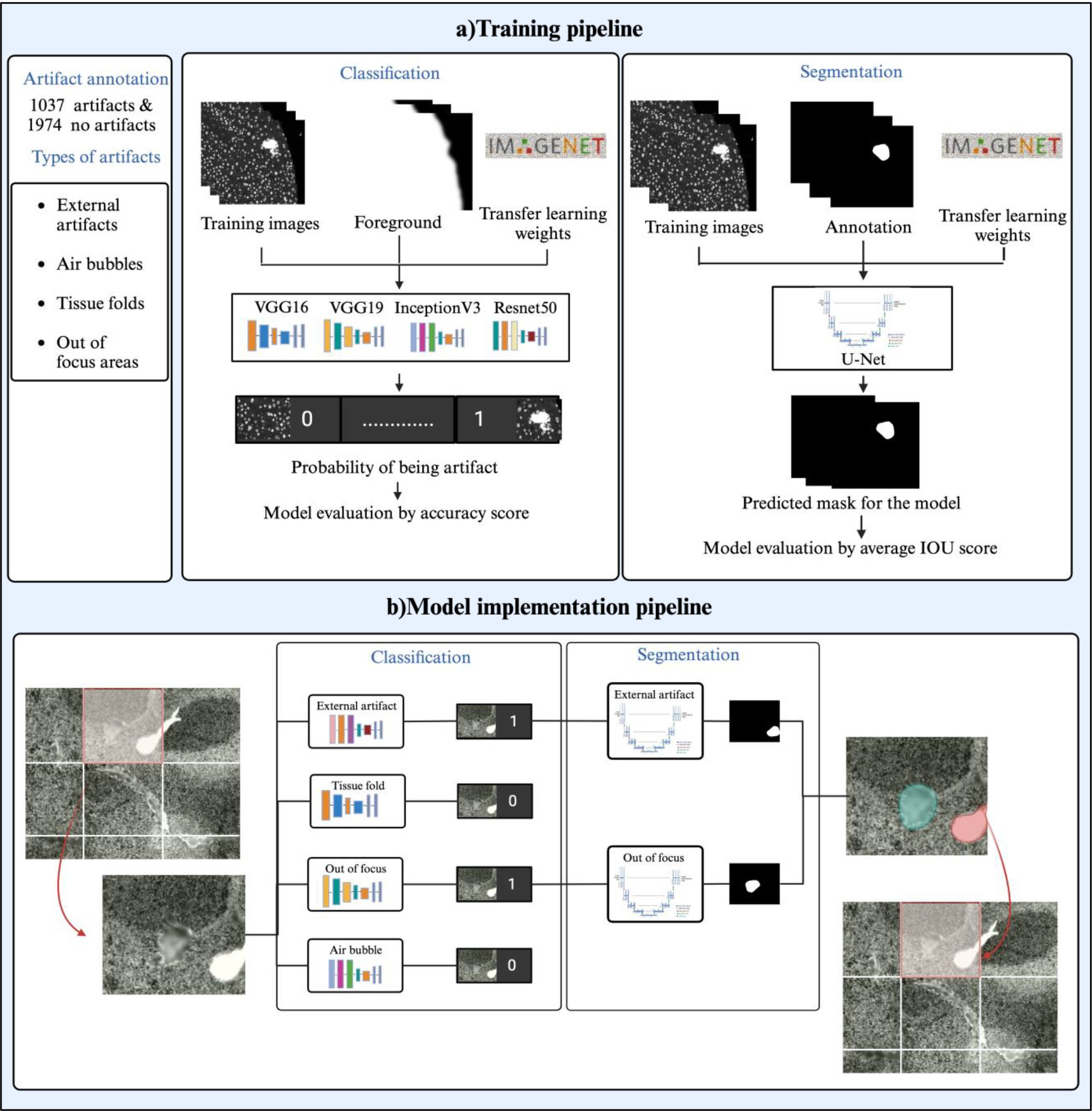
QUAL-IF-AI training and implementation scheme. a) The training pipeline encompasses the pre-processing of the input images as well as the independent training of the classification and segmentation modules for each artifact type. For the training of the classification module, models were trained using transfer learning and four distinct architectures. The loss function of these models was weighted based on the image foreground. The best classification model for each artifact was selected by evaluating accuracy on the validation dataset. For the training of the segmentation model, a U-Net architecture was selected using as a backbone the architecture of the selected classification model in each case. Weights were again transferred from the ImageNet dataset. b) In the model implementation pipeline, each image was divided into tiles of 512 pixels of lateral size, serving as inputs to the previously trained classification models. Tiles identified as having artifacts by the classification model were processed through the trained segmentation model to predict the artifact segmentation masks.

## Materials and methods

### Data collection

A diverse and heterogeneous dataset was built including fully anonymized images from different acquisition technologies: Multiple Iterative Labeling by Antibody Neodeposition – MILAN [24], LUNAPHORE – COMET [25], and AKOYA – CODEX [26]. To increase the generalizability of our models, the collected datasets included a variety of tissue types representing different pathologies, assembled in different types of slides, and acquired at different resolutions. The complete set of parameters considered when collecting these datasets is detailed in Table 1.

**Table 1.** Overview of datasets used for training QUAL-IF-AI models.

| Acquisition technology | MILAN | COMET (LunaPhore) | CODEX (Akoya) |
| --- | --- | --- | --- |
| Species | Human | Human | Human |
| Tissue type | Healthy lung, appendix, brain, liver, and pathologies including metastatic melanoma, pancreatic ductal adenocarcinoma, colorectal cancer, diffuse large B cell lymphoma, autoimmune hepatitis, kidney transplant, glioblastoma | Healthy tonsil, Pan cancer | Metastatic melanoma |
| Slide type | WSI, TMA, CNB | WSI, TMA | WSI |
| Acquisition instrument | Zeiss Axioscan Z1 | LunaPhore COMET | Keyence BZ-X800 |
| Resolution ($\mu\text{m}/\text{px}$) | 0.65 | 0.22 | 0.37 |
| Number of images with artifacts | 995 | 285 | 556 |
| Number of images without artifacts | 1108 | 298 | 620 |

### Image pre-processing and Artifact annotations

The images were processed with zero padding and normalized using q99 normalization. Subsequently images were split into tiles of 512 pixels on each side with a 20-pixel overlap between consecutive slides. For each of the above-mentioned artifacts, 50 randomly selected tiles were visually evaluated and manually annotated by image analysis experts (MDA, TO, JV, MV, BD) using our in-house-built fast labelling tool (see below). The image tiles obtained from the DAPI channel were first categorized based on the presence or absence of artifacts. Annotations were done only on the DAPI channel as it is shown to be sufficient for quality checking in multiplex immunofluorescence images [7]. Further, for images where artifacts were detected, manual artifact segmentation was performed to precisely delineate the areas affected by these artifacts. Following an active learning approach (see below), incrementally more tiles were annotated up to a total of 2,993 (1,019 images containing at least one of the 4 types of artifacts and 1,974 image tiles without any artifact) (Table *1*)

### QUAL-IF-AI fast labeling tool

We have developed a labeling tool to perform a fast annotation of artifacts in the images. This tool is divided in two modules. The first module is the classification module (also called fast-flagging mode), where images from a predefined input directory are sequentially displayed to the user who can flag the images having an artifact by clicking “yes” or “no”. If the user flags an image as having an artifact, then the tool moves forward to the segmentation module (also called labelling mode), where the user can identify the type of artifact and annotate/draw the exact location of the artifact. annotate/draw the exact location of the artifact. Image J plugins are incorporated in the tool to modify the images for better visualization (Fig 2)[27].

**Fig 2.**
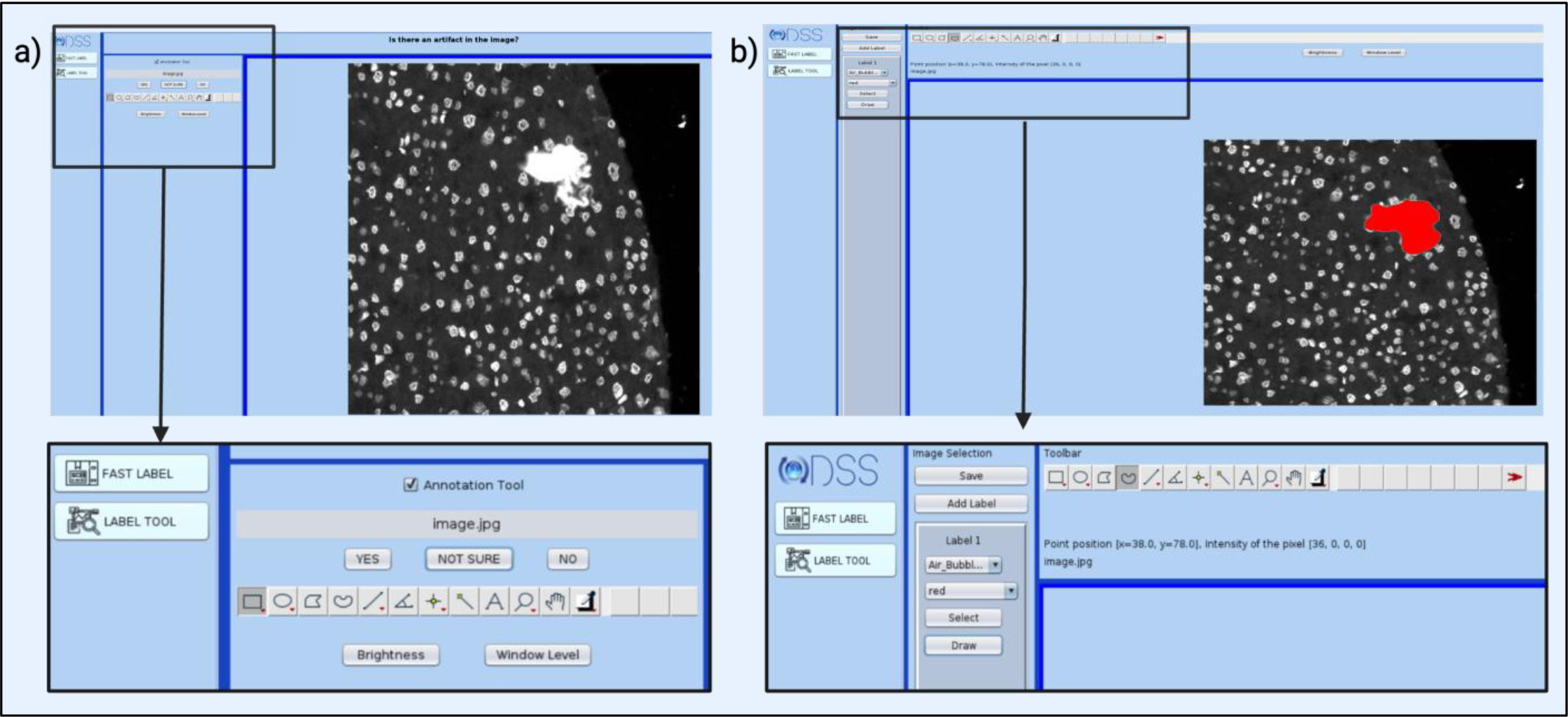
QUAL-IF-AI fast labeling tool. Representative screenshots showing the usage of Qual-IF-AI’s fast labelling tool. A) Flagging mode to classify images with presence/absence of artifacts. B) Labelling mode to identify the type of artifact and annotate the exact location of the artifacts (in red) in the images.

### Model training

Fig 1 summarizes the pipeline followed for model training. Model training was performed independently for each artifact type, and it concatenates a classification and a segmentation module to generate the final output. Tiles were split into training (60%), testing (20%) and validation (20%) datasets. Splitting was performed in a stratified manner keeping a balanced representation of each artifact in each cohort. We started training with 50 image tiles and employed an active learning approach (see below) to selectively add images to the training dataset accelerating model learning. The reader should note that artifact annotations and training were performed exclusively in areas of the image covered by the tissue (tissue foreground) or partially overlapping with the tissue. Therefore, during training, foreground masks were provided together with the annotations of the artifacts. To identify tissue foreground, the ‘COREOGRAPH’ tool from MCMICRO was adopted [4].

#### Classification

For each artifact category, a separate binary classification model was trained to recognize tiles containing a specific artifact. Given that the same tile can have multiple artifacts at the same time, a multi-level classification approach was discarded. Several popular classification architectures were tested for this task. The model architecture and the initial weights were imported from the ImageNet dataset as implemented in the Keras library in python [22]. Using models pretrained on the ImageNet dataset reduces the number of images required for model training [28]. The ‘ImageDataGenerator’ function from Keras was used for data augmentation. The ‘Adam’ optimizer with a learning rate of 0.00001 was used and ‘binary cross entropy’ was selected as loss function. Each model was trained for a maximum of 250 cycles (epochs). The model with highest accuracy on the test dataset was selected by early stopping.

#### Segmentation

For each artifact category, a separate segmentation model was trained to delineate the tissue area covered by the artifact with pixel-level resolution. The U-Net architecture from the ‘segmentation models’ python library was used for this task [23]. The U-Net model consists of two parts, an encoder, and a decoder. The function of the encoder is to extract features and encode the original input image to obtain a fixed-length vector. The function of the decoder is to decode the vector to generate a new output image. The ’
ssegmentation models’ library allows users to choose an encoder architecture with a varying number of parameters [29]. In this study, the model with the highest accuracy in the classification task was used as encoding backbone to build the segmentation model. Weights from the ImageNet dataset were used as a starting point to train the segmentation models. The ‘ImageDataGenerator’ function from Keras was used for data augmentation. The ‘Adam’ optimizer with a learning rate of 0.00001 was used and ‘binary cross entropy’ was selected as loss function. Each model was trained for 200 cycles (epochs).

### Decision Support System for Active Learning

Creating a comprehensive dataset encompassing all possible artifact types, particularly in the first stages of training, is unfeasible. While transfer learning from models trained with general databases such as ImageNet accelerates the process, they are still insufficient for most computer vision tasks. This challenge can be addressed by refining models through the integration of annotated cases from the datasets of interest. In this context, active learning frameworks are increasingly favored over random selection due to several compelling advantages such as training efficiency (they improve model performance with fewer labeled instances) [30], mitigating class imbalance [31] and increasing dataset quality by focusing on ambiguous instances that are most informative for the model [32]. In this line, active learning frameworks heavily rely on Decision Support Systems (DSS) guiding the intelligent selection of the most informative data points to improve model performance efficiently. This targeted approach to data selection leads to a more rapid and effective convergence of the model, particularly in complex tasks such as image classification [33]. After the initial training including 50 tiles, a template was constructed by applying Principal Component Analysis (PCA) on the 128-tensor generated by the dense layer preceding the output layer. Then, new data was projected on this PCA and using a k-nearest neighbors (KNN) (K = 10) approach, these tiles were classified into artifacts (>=8 of 10 nearest neighbors belong to the artifact category), no-artifacts (>=8 of 10 nearest neighbors belong to the no-artifact category), and uncertain (<8 artifacts belong to either category). Then, 50 tiles from the uncertain category were randomly selected and annotated as described above. Finally, the model was retrained, and the process repeated until reaching convergence (Fig 3)

**Fig 3.**
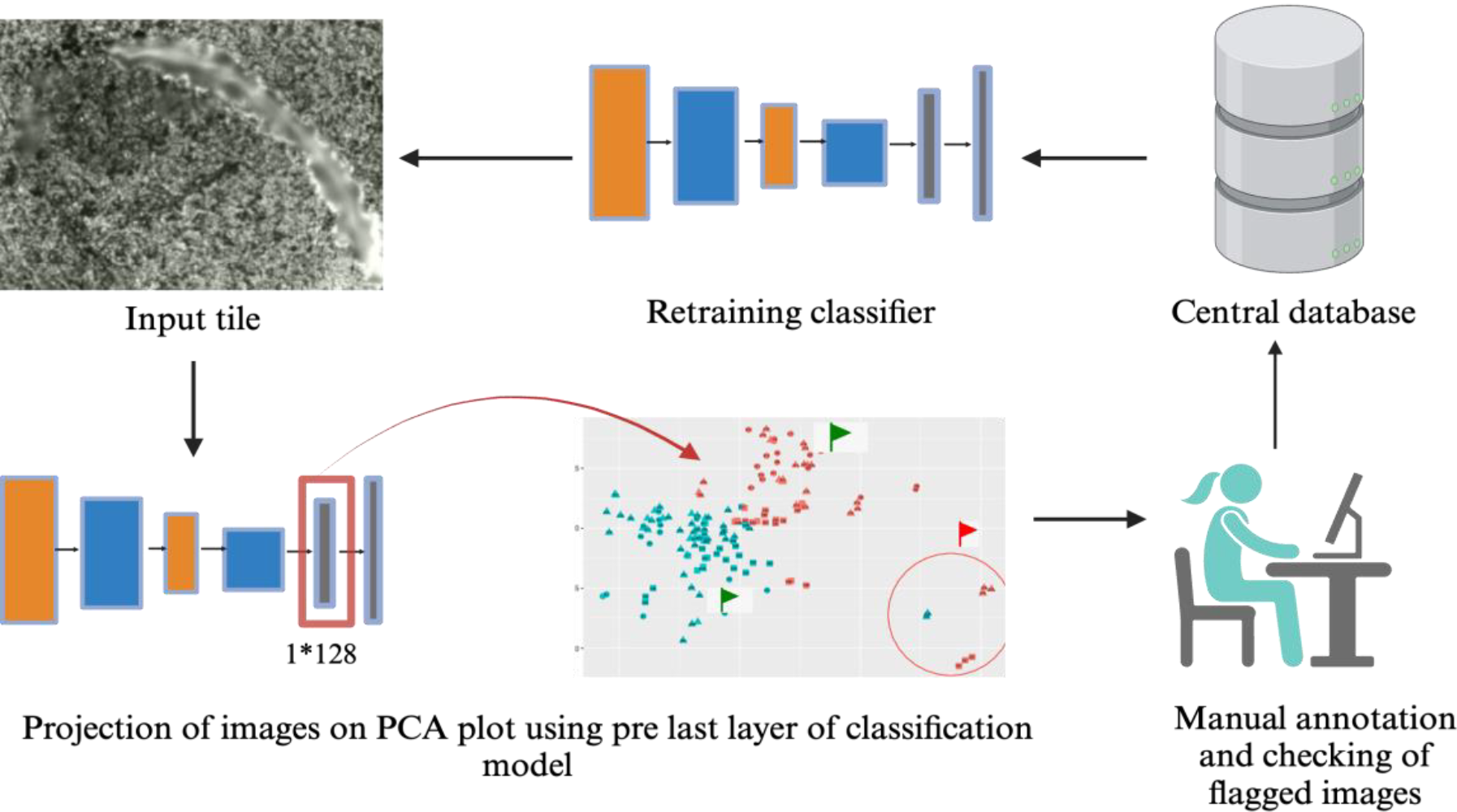
QUAL-IF-AI’s active learning process. PCA plots are generated using the 128-length tensor generated by the dense layer of the classification model. Each new input image is projected on the PCA, and a KNN approach is used to flag the images. Images with at least 8 out of 10 neighbors belonging to the artifact category are classified into the artifact category, whereas images with at least 8 neighbors from non-artifact category are classified into the no-artifact category. Images with less than 8 neighbors of any of the categories are classified as uncertain. Images from the uncertain category are manually annotated and used for training in the next iteration.

### Evaluation metrics

Classification models were evaluated using accuracy, which judges the overall correctness of predictions across all classes and is convenient in a balanced dataset [34]. Accuracy scores were represented together with Received Operating Characteristic (ROC) curves to compare different models for different artifact types. ROC curves provide a graphical representation of the trade-off between the true positive rate and the false positive rate at various threshold values [35].

Segmentation models were evaluated using the Intersection over Union (IoU) metric. IoU is a metric between 0 and 1 that quantifies the overlap between the predicted mask and its corresponding annotation [36]. Generally, an IoU score of 0.5 is acceptable for artifact segmentation [18].

## Results

### Qual-IF-AI is able to identify areas with artifacts

As illustrated in Fig1, the goal of QUAL-IF-AI is to identify areas on fluorescence images affected by different types of artifacts, including OOF, air bubbles, tissue folds, and external artifacts. This is obtained by a two-tier classification and segmentation deep-learning model that first identifies tiles containing a given artifact, and if the prediction is positive, segments the area covered by the artifact with pixel-level precision. To show the performance of QUAL-IF-AI, we collected a series of fluorescent image datasets acquired with different technologies at different resolutions and involving a variety of slide and tissue types (Table1)

QUAL-IF-AI achieved a classification accuracy of over 85% in the validation cohort of all artifacts (Table 2). There were no major differences in accuracy between the different architectures tested for this purpose (Fig 4). For each artifact type, the architecture providing highest accuracy was selected. The Resnet50 architecture achieved a classification accuracy of 91% in classifying out of focus areas and 89% in identifying external artifacts. The Inception V3 model architecture achieved 92% accuracy in classifying tissue folds and 85% accuracy in classifying air bubbles. These architectures were then used as a backbone for their respective segmentation models.

**Table 2.** Details of architectures used in training classification and segmentation models and their performances for each artifact type.

|  | Classification |  |  |  | Segmentation |  |
| --- | --- | --- | --- | --- | --- | --- |
|  | VGG16 | VGG19 | InceptionV3 | ResNet50 | U-Net<br>(ResNet50 encoding) | U-Net<br>(Inceptionv3<br>encoding) |
| **Number of layers** | 16 | 19 | 164 | 50 | 231 | 352 |
| **Total number of parameters** | 138.4 M | 143.7 M | 56M | 23M | 32M | 29M |
| **Accuracy tissue fold** | 0.91 | 0.88 | 0.92 | 0.84 | - | 0.64 |
| **Accuracy air bubble** | 0.84 | 0.81 | 0.85 | 0.80 | - | 0.70 |
| **Accuracy external artifacts** | 0.86 | 0.85 | 0.82 | 0.89 | 0.67 | - |
| **Accuracy OOF** | 0.84 | 0.84 | 0.81 | 0.91 | 0.66 | - |

**Fig 4.**
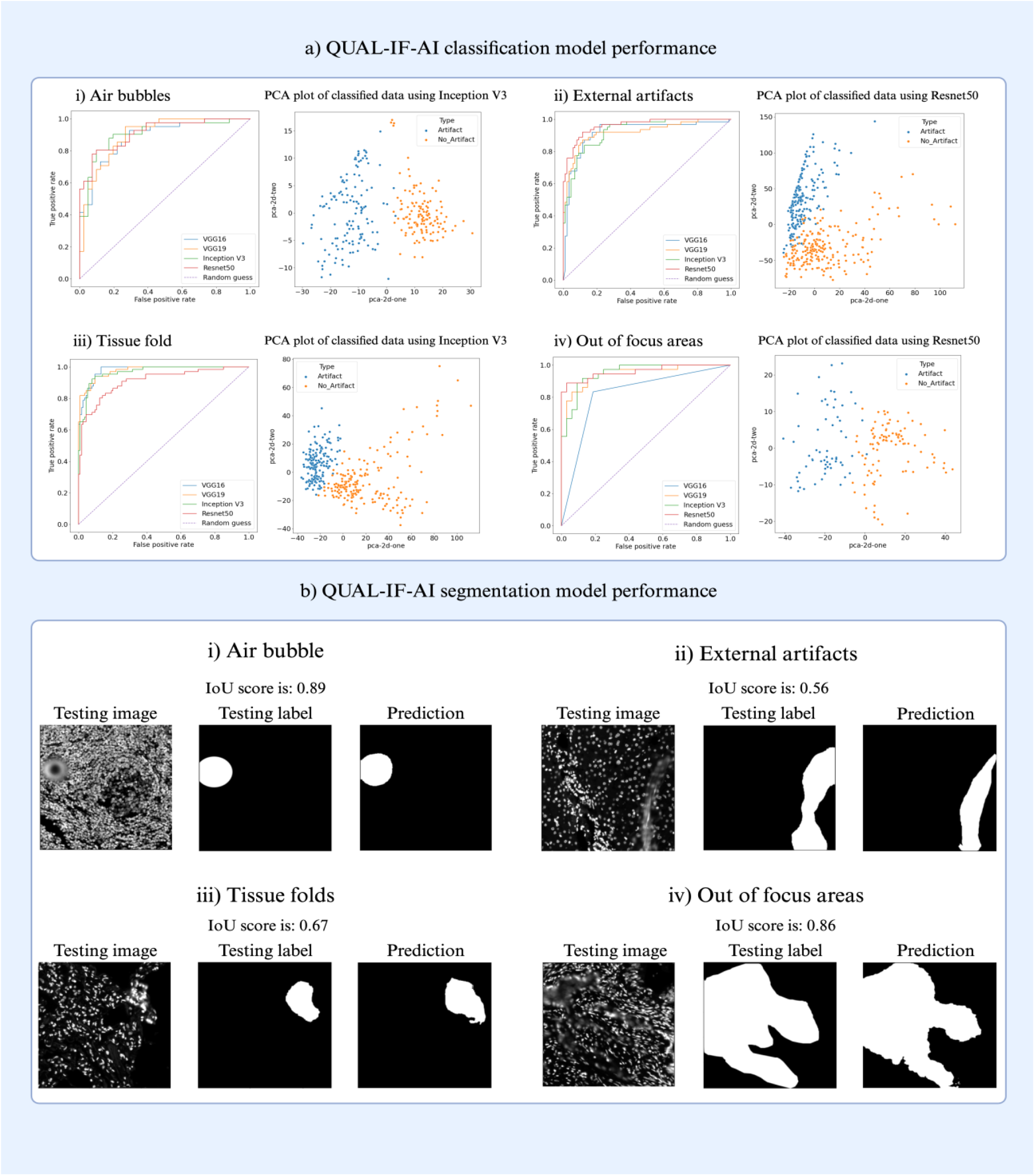
QUAL-IF-AI model performance. a) ROC curves (on the left) and PCA plots (on the right) displaying the performance of classification models for distinct artifacts: (i) Air bubble, (ii) External artifacts, (iii) Tissue fold, and (iv) OOF areas. The ROC curves delineate the model performance, with VGG16 architecture represented in blue, VGG19 in orange, Inception V3 in green, and Resnet50 in red. The purple dashed line denotes the performance of random guessing. In the PCA plot displayed on the left, each dot corresponds to an image from the validation dataset. Images categorized as artifacts are depicted in blue, while those classified as No artifact are represented in orange. b) QUAL-IF-AI segmentation model performance. Segmentation masks as predicted by the segmentation model for specific artifacts: (i) Air bubble, (ii) External artifacts, (iii) Tissue fold, and (iv) OOF regions. Each panel depicts, from left to right, the image containing the respective predicted by the segmentation model on the right.

Image tiles classified as artifacts were further passed through segmentation models. We achieved more than 0.6 IOU (Intersection Over Union) on the validation cohort for segmentation of each artifact type. The model achieved an IoU score of 0.70 for air bubbles, 0.67 for external artifacts, 0.64 for tissue folds and 0.66 for OOF areas. Fig 4 shows an example of the resulting segmentation mask for each artifact category. Fig 5 illustrates an example of the final output of QUALI-IF-AI on a representative image.

**Fig 5.**
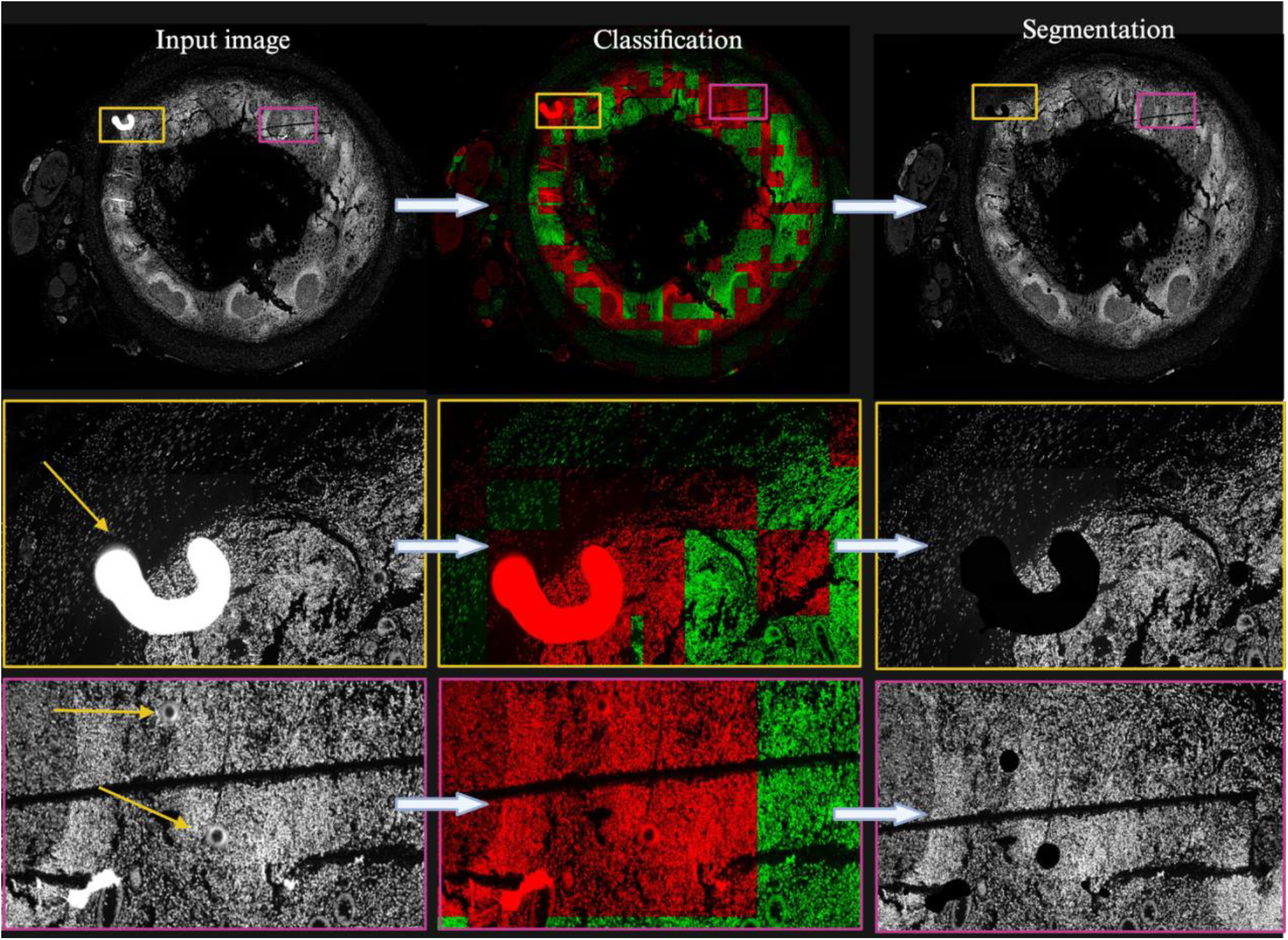
QUAL-IF-AI performance. A representative case of artifact detection. Input images (left) are first passed through the classification module where tiles containing artifacts are flagged (middle) and then the segmentation module removes areas containing artifacts from the images (right). The complete images are shown at the top. Two zoomed in areas from the top images are shown at the bottom. Arrows in yellow indicate artifacts identified in the tissue.

### Improvements from the active learning framework and benchmarking with state of the art

To evaluate the improvement introduced by the active learning framework in comparison with a random tile selection we illustratively selected images with OOF areas and explored the impact of increasing the number of training and testing images on accuracy. Notably, we demonstrated that strategic integration of images, using the active learning methodology as described in the methods, accelerates the model’s increase in accuracy. Through this systematic approach, the model learns faster in comparison to a random selection of tiles. In 10 independent runs, the active. learning approach reaches a mean accuracy of 0.86 after 6 iterations while the random learning reaches a mean accuracy of 0.78 (p-value Wilcoxon test = 1.92e-03) (Fig 6).

**Fig 6.**
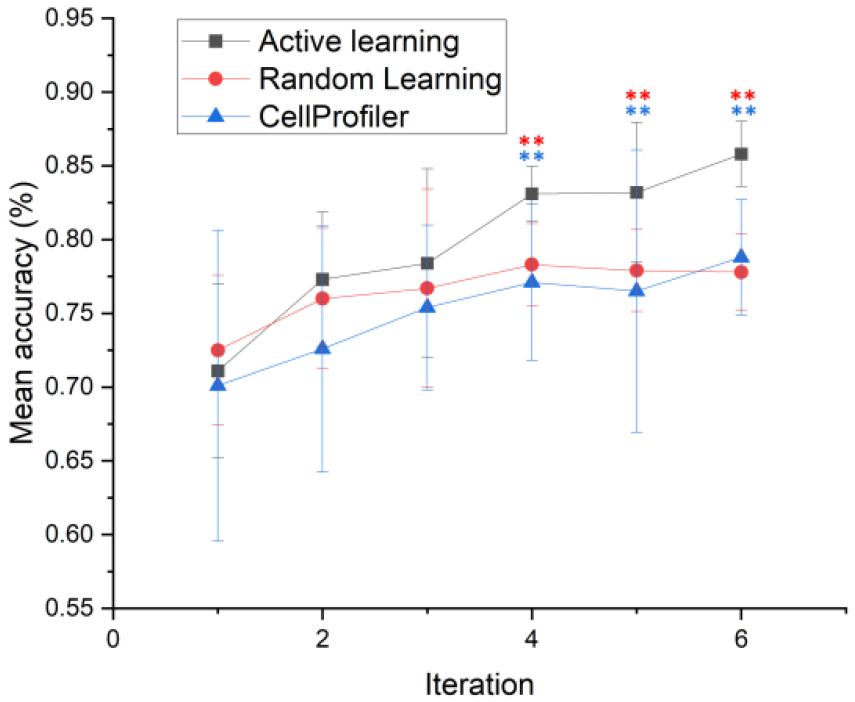
Benchmarking of QUAL-IF-AI with respect to the state-of-the-art. For each method, 10 independent runs were performed, and the accuracy of the classification model evaluated. The black curve depicts the progress under QUAL-IF-AI, the red curve shows the mean accuracy under random sampling of training data and the blue line shows the mean accuracy using CellProfiler tool. Red asterisk 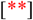 indicate statistically significant difference in the performance of random learning and active learning (p<0.01). Blue asterisk 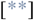 indicate statistically significant difference in the performance of CellProfiler and active learning (p<0.01).

Prior research has demonstrated the potential of utilizing CellProfiler for identifying OOF areas in fluorescent images [16]. In this study, we evaluated the effectiveness of our OOF model against CellProfiler. For this, we performed 6 iterations of training the ‘Random Forest’ classifier with CellProfiler and repeated this 10 times. In this case, the mean accuracy reached by CellProfiler was 0.78 (p-value Wilcoxon test = 8.85e-03) (Fig 6).

## Discussion

Considering the increasing demand for automated quality control in the context of fluorescence microscopy, we developed QUAL-IF-AI, a novel AI-based two-step approach for automated artifact detection and management in fluorescent images. To the best of our knowledge, QUAL-IF-AI represents a first-in-class methodology due to its ability to identify four different common artifacts present in fluorescent images. Our approach ensures a fast and reliable selection of artifact-free images required for high-quality results after image analysis in studies involving immunofluorescence images.

The performance of QUAL-IF-AI was evaluated by using a diverse dataset that was built including images from multiple technologies, tissue types, pathologies, and various groups of artifacts. QUAL-IF-AI focuses on four types of artifacts: Air bubbles, tissue folds, external artifacts and out of focus areas. Nevertheless, the diversity of artifacts present in immunofluorescence images is larger and some less frequent but potentially relevant artefacts such as antibody aggregations have not been included in the method due to their marker dependency. Additionally, QUAL-IF-AI has been evaluated using three different multiplexing technologies. Technologies not including in our study might require fine tuning before the tool provides satisfactory results.

One of the limitations of the study is that it still requires a certain degree of manual intervention for running the models. Therefore, future work will focus on developing a user-friendly software interface to automate and simplify the process further enabling researchers with varying levels of expertise to effectively use our artifact detection method. This enhancement will reduce manual effort and promote widespread adoption among researchers and practitioners in the field.

## Conclusion

In summary, our study presents a first-in-class artifact identification method for immunofluorescence images which achieves a classification accuracy of over 85% and segmentation IoU of more than 0.6 for any of the four artifact types included in the study. We have shown, that adopting an active learning framework accelerates the improvement in accuracy requiring less data annotation. The two-tier classification and segmentation methodology emerge as pivotal for high-quality results reducing the number of false positives. Collectively, our approach advances artifact detection, enabling more accurate image analysis and laying the groundwork for enhanced immunofluorescence staining applications.

## Data availability

The datasets generated and analyzed during the current study are not publicly available at the moment but are available from the corresponding author upon manuscript publication or on reasonable request.

## Acknowledgements

M.D.A. is supported by KU Leuven grant-EHU-D6472-C16/19/006. MV is supported by an FWO PhD fellowship fundamental research (11L0824N). AA is supported by postdoctoral fellowship senior grant from FWO (12ATN24N). FMB is supported by FWO Fundamenteel Klinisch Mandaat EMH-D8972-FKM/20. BDM is funded by KU Leuven Research Fund (projects iBOF/23/064, C16/15/059, C3/19/053, C24/18/022, C3/20/117, C3I-21-00316), KU Leuven Industrial Research Fund, EOS Project no G0F6718N (SeLMA), SBO project S005319N, Infrastructure project I013218N, TBM Project T001919N; PhD Grant (SB/1SA1319N),EWI: the Flanders AI Research Program,VLAIO: CSBO (HBC.2021.0076) Baekeland PhD (HBC.20192204),European Commission: European Research Council under the European Union’s Horizon 2020 research and innovation programme, Foundation ‘Kom op tegen Kanker’, CM (Christelijke Mutualiteit)

